# Real-time visual interactions across the boundary of awareness

**DOI:** 10.1101/237917

**Authors:** Noya Meital-Kfir, Dov Sagi

## Abstract

The relationship between our conscious experiences of the visual world and the neuronal processes involved in the processing of visual information is a central theme of current research. However, it is not yet known whether conscious experiences possess unique quantitative or qualitative processing features. Recently we have shown that two visual stimuli, one consciously experienced and one not, interact as function of features and objects similarity, pointing to preserved visual representations in the absence of perceptual awareness. Here we ask whether these representations can be modified in the absence of perceptual awareness by testing interactions while the unperceived stimulus is being modified outside of one’s awareness. Observers performed a Motion Induced Blindness task, wherein a plaid ‘Target’ was morphed into a Gabor patch once disappearance was reported. Reappearances of the morphed target were induced by a visible ‘Cue’. Our results indicate that the effectiveness of the cue depends on the target representation during the time of suppression. Reappearance rates were highest when the morphed target and the cue share the same orientations. Our findings indicate that the target-cue interactions do not depend on memory-stored representations, but rather on the current state of the consciously unavailable target. We conclude that visual representations can be modified in the absence of conscious perception.

## Introduction

Salient visual stimuli can disappear and reappear from conscious awareness when surrounded by a high-contrast moving background, a phenomenon known as Motion Induced Blindness^1^ (MIB; www.scholarpedia.org/article/Motion_induced_blindness). MIB disappearances are an all-or-none phenomenon and can last up to several seconds, even with a high contrast target located near fixation^2^. It has been proposed that known low-level processes such as contrast adaptation^3^, and filling-in^4^ cannot provide a complete account of MIB, and that a competition between the neural representations of the static target and the moving background needs to be considered in order to explain the range of effects found^1,2,5^.

The all or none nature of MIB disappearances provides a tool to test visual processing at times of perceptual suppression. Indeed, studies have shown that low-level adaptation^6,7^ and grouping by the Gestalt principles^1,8^ are maintained during MIB episodes. In a more recent study^9^ we attempted to specify the information available on perceptually invisible stimuli. Specifically, we tested whether invisible stimuli are represented as a collection of features or as integrated objects^9^. Using a method introduced by Kawabe et al.^10^ we^9^ measured the reappearances of a perceptually invisible ‘Target’ in the presence of a visible ‘Cue’. Interestingly, our results^9^ indicated that the cue interacts with the suppressed target as a function of orientation similarity (~30° bandwidth) and distance (~1° range)^9^. We suggested^9^ that feature-specific interactions between the seen and the unseen imply that information related to the unseen target is available in the system, guiding interactions with the cue. An object-based representation at the subconscious level was tested^9^ by examining the interaction between a compound stimulus, composed of two orthogonal orientations, and its features. Importantly, we showed^9^ that asymmetric relations exist between aware and unaware object representations; a visible plaid cue did not effectively interact with a target defined by only one of its components, but an invisible plaid target efficiently reappeared with component cues and plaid. It was suggested^9^ that perceptually visible as well as invisible objects are represented by combinations of features, possibly at the object level of processing. However, only the perceptually invisible are decomposed into their constituting features^9^.

Our previous findings^9^ indicated similarity-based interactions across the boundary of awareness. However, it is not yet clear whether target-cue interactions during perceptual suppression are affected by the available memory of the target, thus preserving target-related representations available before suppression. To address this issue, we test here the interactions between a visible cue and an invisible target modified during its disappearance, asking whether visual representations that are not consciously available can be modified.

## Methods

### Observers

The experiments include 7 naïve observers (5 females, aged 23–27) with normal or corrected-to-normal vision. Observers gave their written informed consent and were paid for participating. The procedures were in accordance with the Declaration of Helsinki and were approved by the Review Board of the Weizmann Institute.

### Apparatus

The experiments were carried out using the MATLAB Psychophysics toolbox (Psychtoolbox-3; www.psychtoolbox.org;^16^). Stimuli were displayed on a gamma-corrected 23.6’’ VIEWPixx/3D monitor (1920 × 1080, 10bit, 120Hz) viewed at a distance of 100 cm in an otherwise dark room.

### Stimuli

The target was a plaid pattern composed of −45° and +45° Gabor patches (σ=0.12°, ω = 5.67cpd). Each component had a contrast of 50% before its disappearance was reported (see below ‘tasks & procedure’). Cues (50% contrast) were either −45° or +45° Gabor patches, and a plaid pattern composed of the sum of the two orientations (the same as the target). The Target was presented in the upper left quadrant at an eccentricity of 1.5° to a central fixation point (0.31° diameter). The cue was eccentric to the target at a distance of 4*λ* (0.7°) from the target's center. Target and cue were placed within a mask composed of 10×10 black ‘+’ patterns (0.7° width, 1.4° spacing). The complete mask configuration was rotated clockwise, at 2.4 sec/cycle, around the central fixation point. Background-colored “protection zones’’ (0.88° and 0.62° diameter, respectively), around the target and the cue locations, were used to avoid overlap between the stimuli and the mask. Stimuli were displayed over a gray background (mean luminance: 48 cd/m2).

### Tasks & procedures

The Methods used in this experiment were similar to those used in our previous study^9^ with several important modifications. The trial began with the presentation of the MIB display (i.e., target and mask; Figure 1a). The observers fixated on a dot at the display center and pressed a button to report the disappearance of the plaid target. After disappearance was reported, the target was morphed into a Gabor patch (either +45° or −45°) by reducing the contrast of one plaid component (a logistic decay of 150ms). A ‘Cue’ was presented next to the morphed target (150ms delay relative to disappearance), producing nine possible combinations of target and cue (Figure 1b). The morphed target and the cue were presented concurrently for a limited interval of 300, or 400ms before the morphed target was smoothly erased (a logistic decay of 100ms). The Cue was presented until the trial was terminated (800ms total). Observers were instructed to report the number of stimuli they had perceived from the time of disappearance (i.e., zero, one, or two Gabor patches). A report of two perceived stimuli, following suppression, indicated the induced reappearance of the target by the cue. The following trial was initiated by a key press of the observer. The presentation of the cues was randomized within session (30 trials for each morph type × cue × time window combination). For control conditions, each session included trails in which the target remained a plaid pattern (no morph condition) and trials wherein the cue was not presented (spontaneous reappearances).

**Figure 1:**
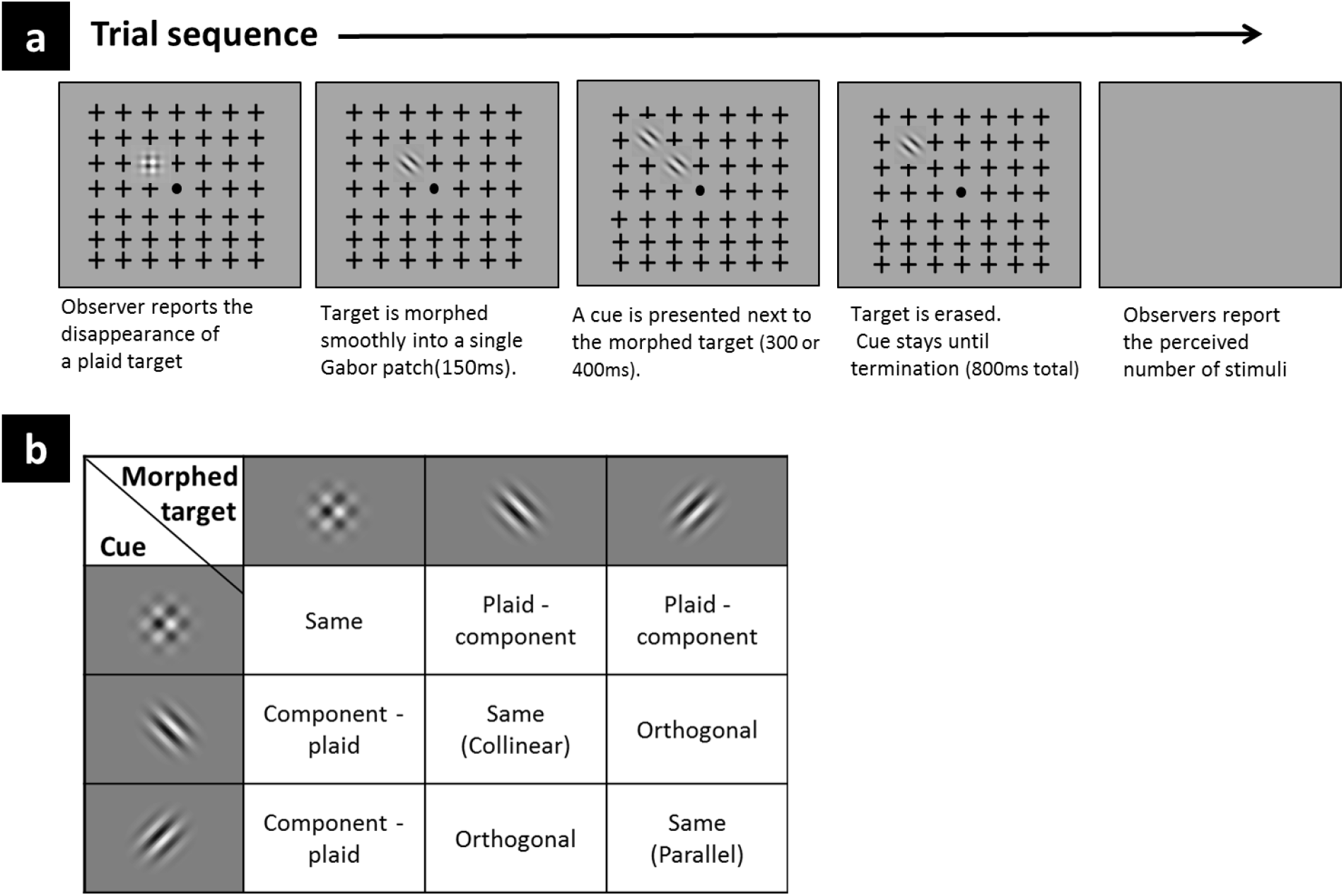
Experimental paradigm; a single trial (**a**). All possible combinations of a morphed target and cue (**b**).

### Data analyses

The frequencies of target reappearances induced by the cue were calculated for each observer as the percentage of trials in which the observer reported two perceived stimuli, from the total number of trials. Repeated-measures ANOVAs were used to test the differences in the frequency of reappearances across conditions. Paired-samples t-tests were used for comparing the differences between two data points.

## Results

To test whether the interactions depend on the representation of the target during the time of suppression, we modified the representation of the unperceived target right after its disappearance. Figure 2 illustrates the frequencies of reappearances as a function of morph and cue type at the two time windows. Rm-anova with two within subject factors (3 × 3) showed a significant morph × cue interaction (*F* (4, 24) = 14.65, *p* = 0.000 and *F* (4, 24) = 16.11, *p* = 0.000, for 300 and 400 ms respectively). Complementary post-hoc analyses indicated that the effectiveness of the cue depends on the target representation during suppression. The frequencies of the reappearances of the morphed Gabor targets were highest in the presence of cues having the same orientation; a −45° morphed target reappeared more frequently in the presence of a collinear cue (−45°) than in the presence of a plaid cue (300ms: *t*(6) = 2.63, *p* = 0.03; 400ms: *t*(6) = 6.67, *p* = 0.001), whereas for the +45° morphed target a parallel cue (+45°) induced higher frequencies of reappearances relative to a plaid cue (300ms: *t*(6) = 3.68, *p* = 0.01; 400ms: *t*(6) = 2.43, *p* = 0.05). Furthermore, there were no significant differences between orthogonal cues and a plaid cue or between a plaid cue and no cue trials (i.e., spontaneous reappearances) for both morphed Gabor targets.

**Figure 2:**
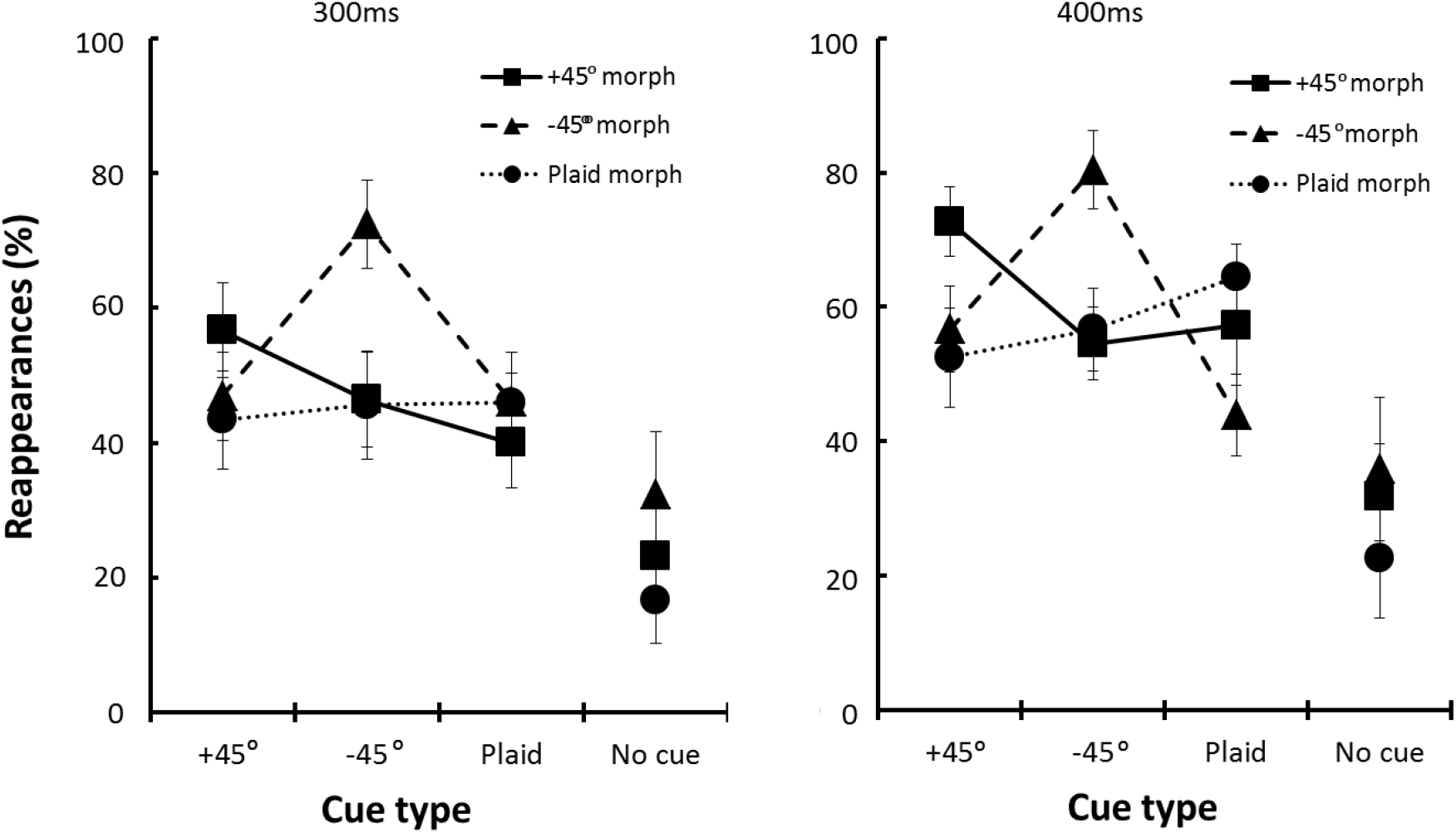
The frequencies of target reappearances as a function of morph and cue type at the 300ms and 400ms time windows. Data points represent means across observers (N=7), error bars represent the standard error of the mean.

In line with our previous finding, indicating that unperceived objects are represented at multiple levels of abstraction, the reappearances of a morphed plaid target were not significantly different between cues.

To control for a possible effect of spontaneous reappearance, the results were converted into d’^11^, cue-driven reappearances were considered as Hits and spontaneous reappearances as False-Alarms (by subtracting the z scores of the induced reappearances from the z scores of spontaneous reappearances). This analysis showed a pattern of results (Figure 3) similar to that observed with the frequencies of reappearances, despite a larger inter-observer variability in cue sensitivity (as indicated by the larger SEs in Figure 3). An exception is the results of the 400ms time window with the +45° morphed target, showing only a marginally significant difference (p<0.06) between a parallel cue (+45°) and a plaid cue. The d’ analysis, by capturing the difference between cue-driven reappearances and spontaneous reappearances, revealed a relatively high sensitivity to the plaid target despite the lack of cue selectivity in agreement with our previous results^9^.

**Figure 3:**
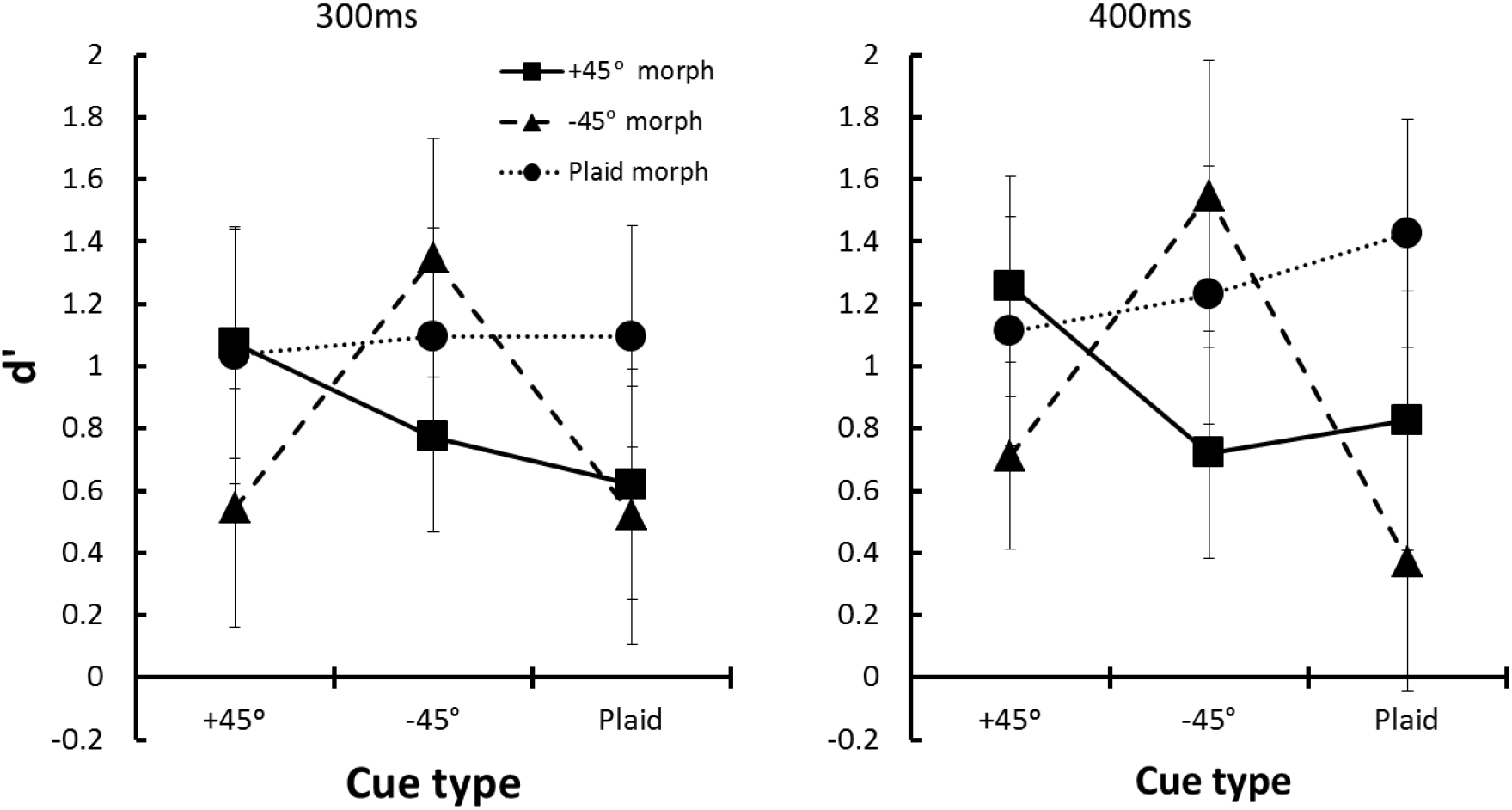
d’ results as a function of morph and cue type at the 300ms and 400ms time windows. Data points represent means across observers (N=7), error bars represent the standard error of the mean.

## Discussion

In the current work we used the MIB paradigm to test the reappearance of perceptually invisible stimuli upon the presentation of a visible cue^10^. After observing similarity-dependent interactions between the aware and the unaware^9^, we were encouraged to ask whether the interactions depend on the representation of the invisible target during the time of suppression and whether these representations can be modified in the absence of visual awareness. We suggested that selective interactions between the modified target representation and the cue imply that visual representations are continuously processed and can be updated during perceptual suppression. Here a plaid target was morphed into a Gabor patch, once the observers reported its disappearance. Overall, our findings indicate real-time interactions between the perceptually visible and the perceptually invisible stimuli, demonstrating that visual representations can be modified even in the absence of conscious perception. The cues induced the reappearances of the target depending on its representation during perceptual suppression; a morphed Gabor target reappeared more frequently in the presence of a Gabor cue with the same orientation than with a plaid cue, the latter exhibited behavior that was not significantly different than spontaneous reappearances. These selective interactions, between the cue and the modified representation of the target, indicate high sensitivity to unperceived changes in visual representations even without conscious perception. Furthermore, the results suggest that the interactions do not depend on a past representation, as perceived before perceptual suppression, but instead indicate ongoing visual processes across awareness states.

The results showed that an invisible plaid target efficiently and uniformly reappeared with component cues and a plaid cue, but that a plaid cue produced target-specific reappearances. The current results converge with our previous findings in showing non-symmetric interactions between the perceived and the unperceived stimuli^9^; an invisible object target was induced to reappear by its visible component cues and the object cue, however a visible object cue was not effective with component targets. It was proposed that out of awareness a plaid is represented at different levels of abstractions, including both features and objects. Therefore, we suggested that subconscious representations include objects and their components, whereas conscious perception is confined to object representations.

The involvement of conscious perception in the formation of visual representations is in the focus of research on visual awareness^12–15^. In a MIB study, Mitroff & Scholl^8^ showed that perceptual groups can be formed outside of awareness, through spatial integration. Changes made in the grouping relations of unperceived stimuli, during MIB suppression, affect their simultaneous reappearance. In the current work we changed the properties of a single stimulus, decomposing objects into a single component. Moreover, here the representations of the invisible were tested implicitly, by showing interactions across awareness states between an invisible stimulus and its visible context.

In the current study we show a real-time interaction between a perceptually suppressed stimulus and its visible context. By changing the properties of an unperceived object during perceptual suppression, we showed a similarity-dependent interaction between the visible stimulus and the modified representation of the invisible stimulus. Our findings show high sensitivity to unperceived changes in representation, indicating that visual representations can be modified without conscious perception.

## Acknowledgements

We thank Drs. Yoav kfir and Michail Katkov for assisting with the technical aspects of the research, Dr. Tomer Livne for helpful discussions and comments.

This research was supported by a grant from the US-Israel Binational Science Foundation (BSF-2007224), and by The Weizmann Braginsky Center for the Interface between the Sciences and the Humanities.

## Affiliations

Department of Neurobiology, Weizmann Institute of Science, Rehovot 76100, Israel

## Contributions

N.M-K. and D.S. designed the study; N.M-K. performed the experiments and analyzed the data; N.M-K. and D.S. wrote the paper.

## Competing interests

The authors declare no competing financial interests.

## Corresponding author

Correspondence to

